# Preprocessing Choices for P3 Analyses with Mobile EEG: A Systematic Literature Review and Interactive Exploration

**DOI:** 10.1101/2024.04.30.591874

**Authors:** Nadine S. J. Jacobsen, Daniel Kristanto, Suong Welp, Yusuf Cosku Inceler, Stefan Debener

**Affiliations:** Neuropsychology Lab, Department of Psychology, Carl von Ossietzky University Oldenburg, Germany; Psychological Methods and Statistics Division, Department of Psychology, Carl von Ossietzky Universität Oldenburg, Germany; Cluster of Excellence Hearing4all, Carl von Ossietzky Universität Oldenburg, Oldenburg, Germany; Centre for Neurosensory Science & Systems, Carl von Ossietzky Universität Oldenburg, Oldenburg, Germany

**Keywords:** electroencephalography, EEG, mobile, preprocessing, systematic literature review, shiny app

## Abstract

Preprocessing is necessary to extract meaningful results from electroencephalography (EEG) data. With many possible preprocessing choices, their impact on outcomes is fundamental. While previous studies have explored the effects of preprocessing on stationary EEG data, this research delves into mobile EEG, where complex processing is necessary to address motion artifacts. Specifically, we describe the preprocessing choices studies reported for analyzing the P3 event-related potential (ERP) during walking and standing. A systematic review of 258 studies of the P3 during walking, identified 27 studies meeting the inclusion criteria. Two independent coders extracted preprocessing choices reported in each study. Analysis of preprocessing choices revealed commonalities and differences, such as the widespread use of offline filters but limited application of line noise correction (3 of 27 studies). Notably, 59% of studies involved manual processing steps, and 56% omitted reporting critical parameters for at least one step. All studies employed unique preprocessing strategies. These findings align with stationary EEG preprocessing results, emphasizing the necessity for standardized reporting in mobile EEG research. We implemented an interactive visualization tool (Shiny app) to aid the exploration of the preprocessing landscape. The app allows users to structure the literature regarding different processing steps, enter planned processing methods, and compare them with the literature. The app could be utilized to examine how these choices impact P3 results and understand the robustness of various processing options. We hope to increase awareness regarding the potential influence of preprocessing decisions and advocate for comprehensive reporting standards to foster reproducibility in mobile EEG research.

## Preprocessing Choices for P3 Analyses with Mobile EEG: a Systematic Literature Review

Electroencephalography (EEG) is a popular and versatile method for capturing brain-electrical activity. Mobile EEG has gained some momentum over the past years, as it allows the capture of neural activity from participants during whole-body motion (Debener et al., 2012; Niso et al., 2022; Wascher et al., 2021) and promises the investigation of cognition in natural settings. Yet, to fulfill this potential, adequate artifact processing is paramount. Whenever EEG signals are acquired, neural activity is captured along with activity from various other physiological (e.g., heartbeats, eye blinks, and movement; muscular activity from the scalp, face, neck, and shoulders) and non-physiological sources (wire and electrode motion, high impedances, line noise). Disambiguation of neural from non-neural signal contributions, which is the goal of EEG preprocessing, is essential (Thompson et al., 2008). Since EEG artifacts can dominate the recordings during participant motion, applying appropriate preprocessing is even more important when analyzing the EEG of moving participants (Jacobsen et al., 2021; Klug & Gramann, 2020; Nordin et al., 2020).

Various preprocessing steps and parameters could be used, so researchers face a vast number of decisions and degrees of freedom when deciding on their preprocessing pipeline. They not only have to decide which steps to use but also with which parameters, and in which order. These choices lead to multiple branching paths, creating a so-called *garden of forking paths* (Gelman & Loken, 2013). A large variability in preprocessing choices may increase the variability of results and contribute to replication difficulties (Simmons et al., 2011; Trübutschek et al., 2024). An interesting way to tackle this problem is to investigate the impact of analytical decisions on study results. This approach has been termed multiverse analysis (Steegen et al., 2016). Multiverse analyses, for instance, compare the impact of previously identified processing pipelines, thereby reducing the problem of selective reporting and increasing transparency and robustness. Multiverse analyses depend on a proper identification of defensible, that is justifiable, forking paths. While novel approaches allow the investigation of large multiverses (Dafflon et al., 2022), the inclusion of not defensible choices may bias results (Del Giudice & Gangestad, 2021). The processing steps and combinations included in a multiverse analysis may be chosen by the researcher a priori (Clayson et al., 2021; Sadus et al., 2023; Schubert et al., 2023), empirically sampled with a many analysts approach (Botvinik-Nezer et al., 2019; Trübutschek et al., 2022) or determined by a literature review (Šoškić et al., 2021). For instance, a systematic literature review of 132 stationary EEG publications revealed that no two publications chose the same approach of data recording, processing, and analysis and that most omitted at least some details – even while they all examined the same event-related potentials (ERPs) (Šoškić et al., 2021). A multiverse analysis of a subset of the identified preprocessing choices showed that these deviations influence the obtained ERP (Šoškić et al., 2022). Moreover, robust preprocessing depends on the investigated ERP component (Clayson et al., 2021). Yet so far this has not been investigated for ERPs recorded with mobile EEG, although mobile capture of brain activity is a prerequisite for understanding neural activity in more ecologically valid settings and is becoming increasingly popular (Gramann, 2024; Niso et al., 2022). The P3 is an ERP component extensively studied during whole-body motions like walking. Changes in the P3 (i.e., decreased amplitude and increased latency) were repeatedly reported and have been attributed to diversion and re-allocation of cognitive resources such as attention toward motor control (De Vos, Gandras, et al., 2014; De Vos, Kroesen, et al., 2014; Debener et al., 2012; Ladouce et al., 2019).

Here, we provide a systematic literature review to identify and document existing mobile EEG preprocessing strategies of the P3 during walking and standing. The definition of defensible forking paths is a prerequisite for a multiverse analysis of preprocessing choices to systematically assess their impact. Moreover, we introduce an interactive visualization, implemented as a Shiny app (Chang et al., 2023), that allows users to evaluate previously used mobile EEG preprocessing choices and explore how frequently particular steps, and combinations thereof, are used. This application is an adaptation of previous work illustrating the processing of functional magnetic resonance imaging (Kristanto et al., 2024). The app is designed to identify whether a planned preprocessing pipeline has been used before or requires additional justification and validation. The app allows for the easy identification of common, debated, and unique preprocessing choices, including an exemplary preprocessing pipeline that will help researchers new to mobile EEG analysis to navigate safely through a rich garden of forking paths.

## Methods

### Literature search

We followed the preferred reporting items for systematic reviews and meta-analyses (PRISMA) guidelines (Moher et al., 2009) (see supplemental for checklist) and searched the literature using three databases (Scopus, Web of Science, and PubMed) for publications containing the terms *EEG or ERP and P3 or P300 and gait*. Default settings for search engines were used, including the search for terms in the title, abstract, and keywords for Scopus, the search within *topic* for the Web of Science, and the search for *Medical Subject Headings* in PubMed. The search was conducted on June 7, 2022 (Scopus, Web of Science) and on September 26, 2022 (PubMed). All search terms and the respective date of the search, as well as the number of results, are available in the supplemental materials. All references were merged into one database using Zotero (Corporation for Digital Scholarship, Vienna, VI; RRID:SCR_013784) and duplicates were removed with the built-in functionality.

### Study Inclusion

All abstracts were screened by two independent reviewers (NSJJ, SW) to determine whether the publication could be included. Only studies meeting all of the following criteria were included: (1) empirical study (no reviews or book chapters), (2) EEG data were recorded, (3) a P3 was analyzed,(4) the P3 was analyzed during a gait condition (treadmill, overground, exoskeleton), (5) the P3 was analyzed during a non-movement condition, such as sitting or standing, (6) the study was performed on humans, (7) EEG data of healthy participants were analyzed. If the necessary information could not be obtained from the abstract, the reviewers checked the remaining sections of the article.

We only included studies in which both the P3 during walking and in a stationary condition (e.g., standing, sitting) were analyzed. This was a prerequisite as we targeted the P3 alteration with motion. Moreover, we only included healthy participants as some clinical groups may prompt an adapted EEG preprocessing (e.g., due to altered muscular activity). In cases where a patient population was investigated, the presence of a healthy control group for which the P3 was analyzed, was sufficient for study inclusion. Agreement of study inclusion was assessed in percent. In cases of disagreement, a third independent reviewer (SD) was asked, and all three reviewers together agreed on including or excluding the study.

### Coding of preprocessing pipelines

All data was coded by two independent reviewers in separate spreadsheets. Discrepancies were resolved by both reviewers together. Information on the recording setup and task (here called *EEG metadata*) and the preprocessing were extracted. Information on the parametrization of the P3 (e.g., time window, channel(s)) was also coded but is not reported in this publication. If information could not be located within the text of a publication it was coded *NA* and is displayed as *not reported* within the Shiny app.

#### EEG metadata

Among others, data were extracted on the following properties: the task participants performed (e.g., an auditory oddball), power line frequency (in Hz), the manufacturer model, the number of recorded EEG channels, the location of the online reference electrode, the EEG sampling frequency (in Hz), the cut-off of the online filters, the electrode type (dry or wet, active pre-amplification at the recording site or passive transmission), and the employed analysis software. If a study used several systems (for instance to evaluate the performance of different systems), details of all systems were coded. It was noted which open scholarship practices, i.e., open code, open data, or open materials, were employed.

#### Preprocessing

The *Agreed Reporting Template for EEG Methodology - International Standard: template for event-related potentials (ERP) (*ARTEM-IS) (Šoškić et al., 2023; Styles et al., 2021) was adapted to code the EEG preprocessing. The template was extended with the groups *data decomposition* and *source estimation* while coding the latest 10 publications and used to code the remaining 17 studies. The resulting scheme was presented to experienced (mobile) EEG users who recommended including further preprocessing steps for the construction of user pipelines, namely *adaptive filtering*, *canonical correlation analysis (CCA)*, *beamformer*, and *distributed source estimation.* If no specific information about the order of preprocessing steps was provided in the text (“Before we did step 2, we performed step 1”), it was assumed that processing steps were performed in the order they were mentioned in the text. All steps were sorted into the following groups: offline filter, downsampling, re-referencing, reject data, reject channels, interpolate channels, multi-step automated approach, data decomposition, source estimation, line noise correction, epoching, baseline correction, other. All steps are listed in the Shiny app under *Database>List of steps*. Most groups comprise several steps. The offline filter group, for instance, includes low-pass, high-pass, bandpass, and notch filters. Each step is characterized by an option. In the case of offline filters, this was the frequency cut-off in Hertz. The available options were incrementally added while coding all studies. An overview of all available options can be found in the Shiny app *Database>List of Options*. Generally, only the preprocessing for the P3 analysis was coded. If other analyses requiring different preprocessing steps (e.g., analysis of oscillatory activity, auditory attention tracking) were conducted, these were not coded. The same holds for the preprocessing of non-EEG data. For instance, preprocessing of motion sensor data, such as acceleration for gait detection, was omitted (for the use of motion sensors in artifact detection, see e.g.) (Jacobsen et al., 2021). Moreover, preprocessing for data visualization only (e.g., additional low-pass filters) was not extracted. Specific preprocessing targeted at clinical groups (but not controls) was omitted.

### Analysis

Inter-rater reliability of study inclusion was assessed as the percentage of the studies that were included or excluded by both reviewers compared to the total number of studies. Descriptive statistics are available in a Shiny app. In the results section we will summarize our findings by posing and answering questions possible (novice) mobile EEG users may ask themselves.

## Results

### Study Inclusion

All in all, 288 studies were identified of which n = 130 were duplicates. Of the remaining n = 158 studies n = 27 satisfied all inclusion criteria and were analyzed in this study (see Table 1). Both reviewers included the same studies and reached a consensus of 100%. Details on the reasons for study exclusion are detailed in Figure 1.

**Figure 1.**
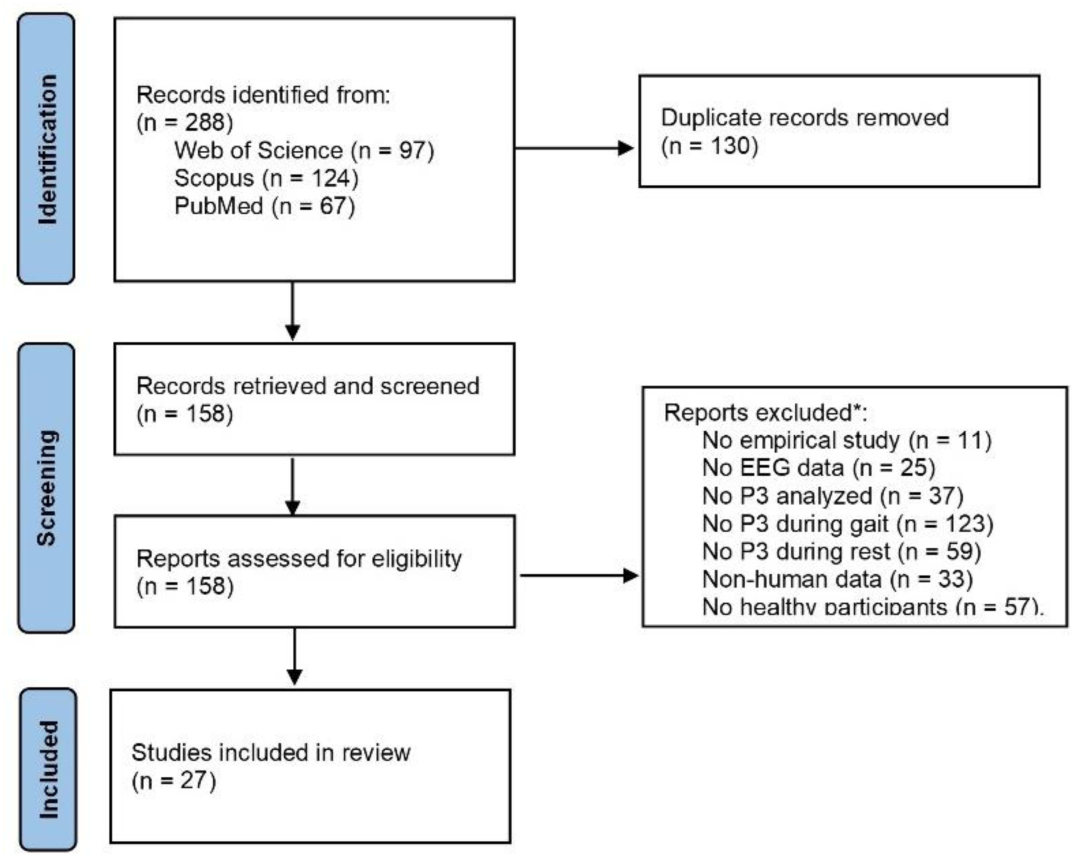
PRISMA-flow diagram for study inclusion. * Reasons for exclusion as coded by reviewer 2. Some studies were excluded for multiple reasons.

**Table 1.**
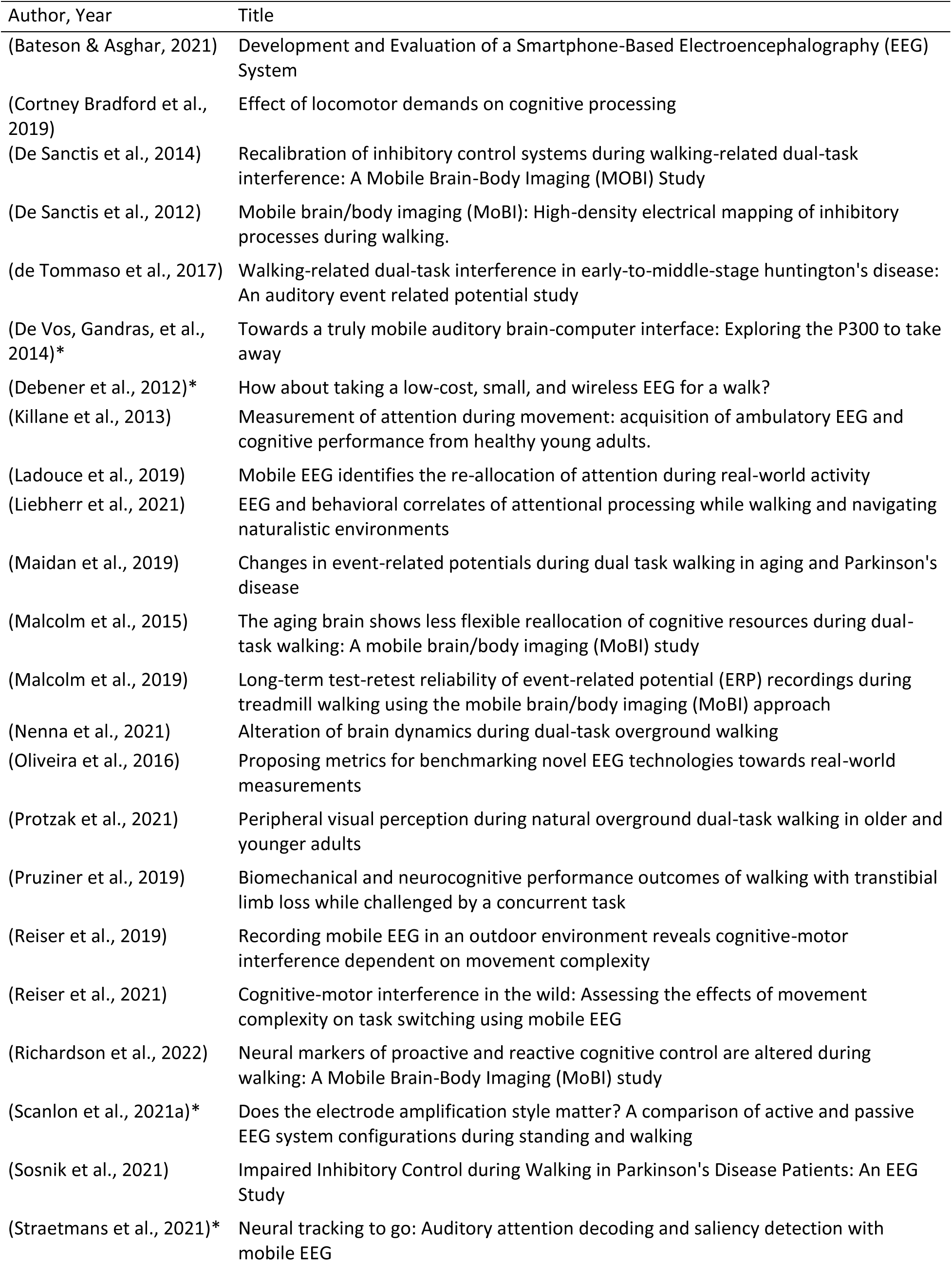
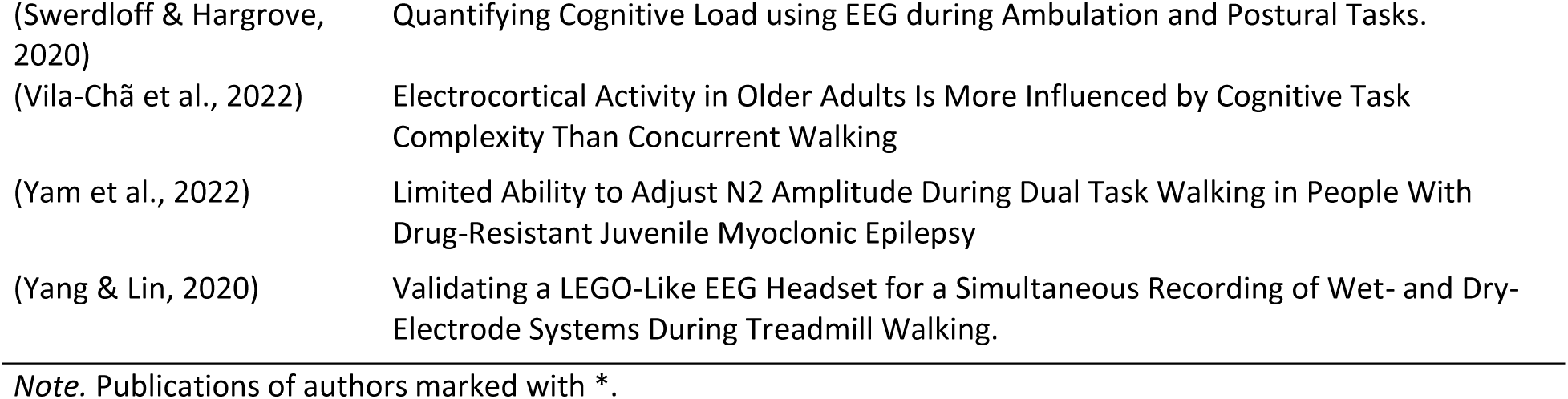
List of included studies.

### EEG preprocessing

On average articles reported 9.67 processing steps (*SD* = 3.69, range 3-16). Each article described a unique preprocessing approach. In total 59,23% of the articles described using at least one manual-preprocessing step (*M* = 1.01, *SD* = 1.11, range 0-4), such as manual rejection of continuous data, epochs, or independent component (IC) selection. For each employed preprocessing step, the chosen option should be reported in an article. Yet at least one option was missing from 55.56% of the articles (*M* = 0.74, *SD* = 0.82, range 0-3).

Steps used by most of the publications include epoching (*n* = 27), baseline correction by subtraction of the mean (*n* = 22), ICA decomposition (*n* = 20), re-referencing (*n* = 19), and bandpass filtering (*n* = 17). Very few studies reported using, dipole fitting (*n* = 3), line noise correction (*n* = 3), or PCA-based methods (n=3).

Details on steps, such as baseline correction (start and end time in ms), bandpass filtering (filter cut-offs in Hz) or re-referencing (name of the electrode) were reported by all studies. Most often omitted details (i.e., not reporting the option for a certain step) are the ICA algorithm (*n* = 6, 30%), identification of correct responses (*n* = 5, 63%), and the method of channel interpolation (*n* = 3, 33%).

### Open Scholarship

Open scholarship practices such as the sharing of analysis code (open code), the sharing of acquired data (open data) or of experimental stimuli and apparatus (open methodology) are not widely used. Only authors of one study shared their code and materials, while authors of four studies shared their data on a repository and another four would share it personally upon request. In several cases (code: *n* = 2, data: *n* = 3) authors intended to share but the links were missing from the publication (see Figure 2).

**Figure 2.**
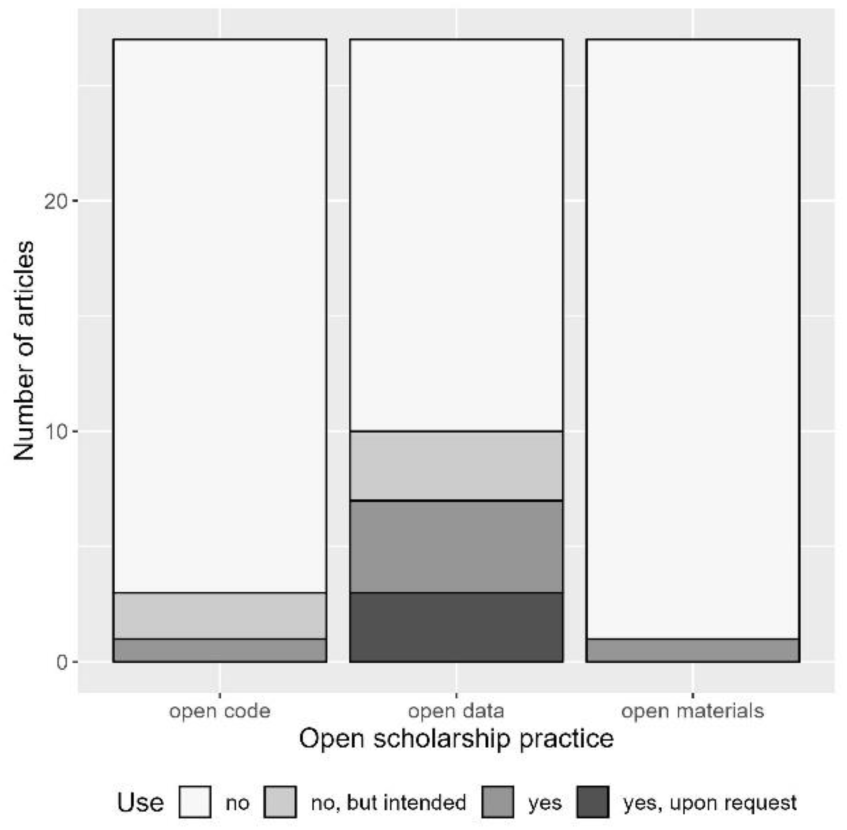
Number of studies using open scholarship practices (open code, open data, open materials).

### Data exploration with the Shiny app

In the following, we compiled a list of possible questions that (novice) EEG users may ask themselves when they set up a new preprocessing pipeline. The Shiny app can be used to address open questions. Instructions on how to find relevant information within the app are indicated in Table 2.

**Table 2.**
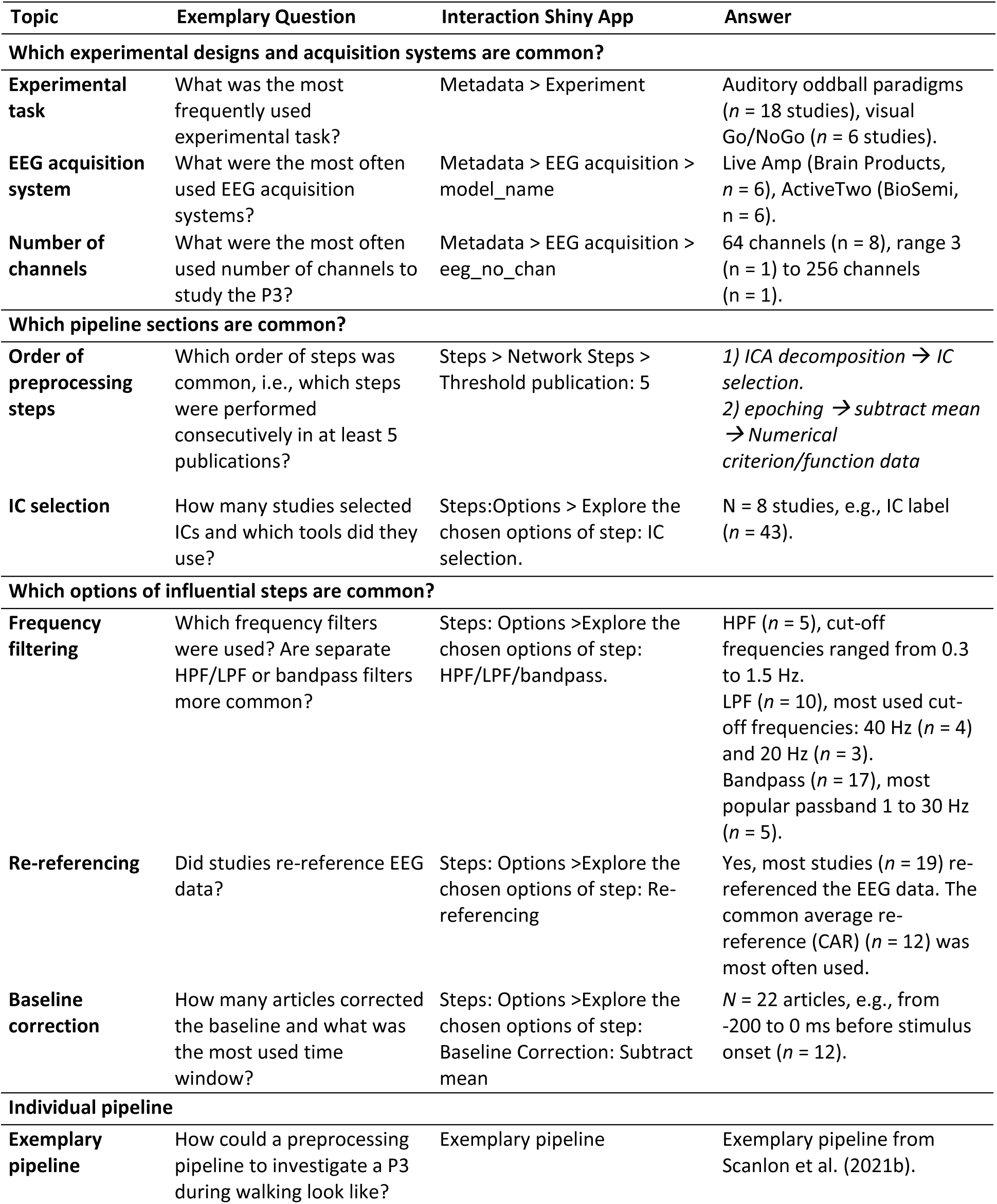
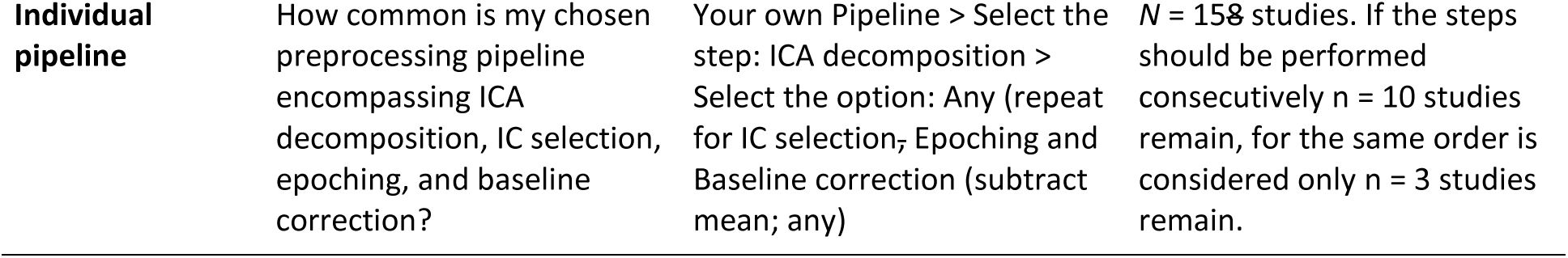
Instructions on how to find relevant information within the shiny app.

#### Which experimental designs and EEG acquisition systems are common?

To explore the P3 effect, most studies used an auditory oddball (*n* = 18 articles) or visual Go/NoGo (*n* = 6 articles) paradigm. EEG data was mostly captured using the Live Amp (Brain Products GmbH, Gilching, GER) (*n* = 6 systems^1^) and the ActiveTwo (BioSemi, Amsterdam, NL) *(n* = 6 systems). The Live Amp is a portable system, designed to measure the EEG of moving participants while the Active Two is designed as a stationary system but can be placed next to participants walking on a treadmill. The most common number of EEG channels was 64 (*n* = 8 systems) while the P3 effect was captured with as few as 3 channels (*n* = 1 system) and up to 256 (*n* = 1 system).

#### Which pipeline sections are common?

While no two pipelines were the same, some steps were often performed in the same order. There were in total four edges (i.e., steps performed in succession) that were used by at least five publications (see Figure 3). These were *ICA decomposition* to *IC selection.* And from *epoching* to subtract mean (*baseline correction) to Numerical criterion/function (data rejection) as well as from epoching to Numerical criterion/function (data rejection)*. Only seven studies did not employ ICA. ICA may yield ambiguous components and component selection can be a source of variation (Pion-Tonachini et al., 2019). Most studies reported using manual (n = 11) or automatic (n = 5) IC selection. Among the automatic tools, IC label (Pion-Tonachini et al., 2019) (*n* = 3) was most frequently used. Other used automatic tools include SASICA (Gramann et al., 2010)(*n* = 1), ADJUST (Mognon et al., 2011)(*n* = 1), selection by the Brain Vision Analyzer (Brain Products, Gilching, GER) and information from dipole fitting (dipole location and explained variance, *n* = 2).

**Figure 3.**
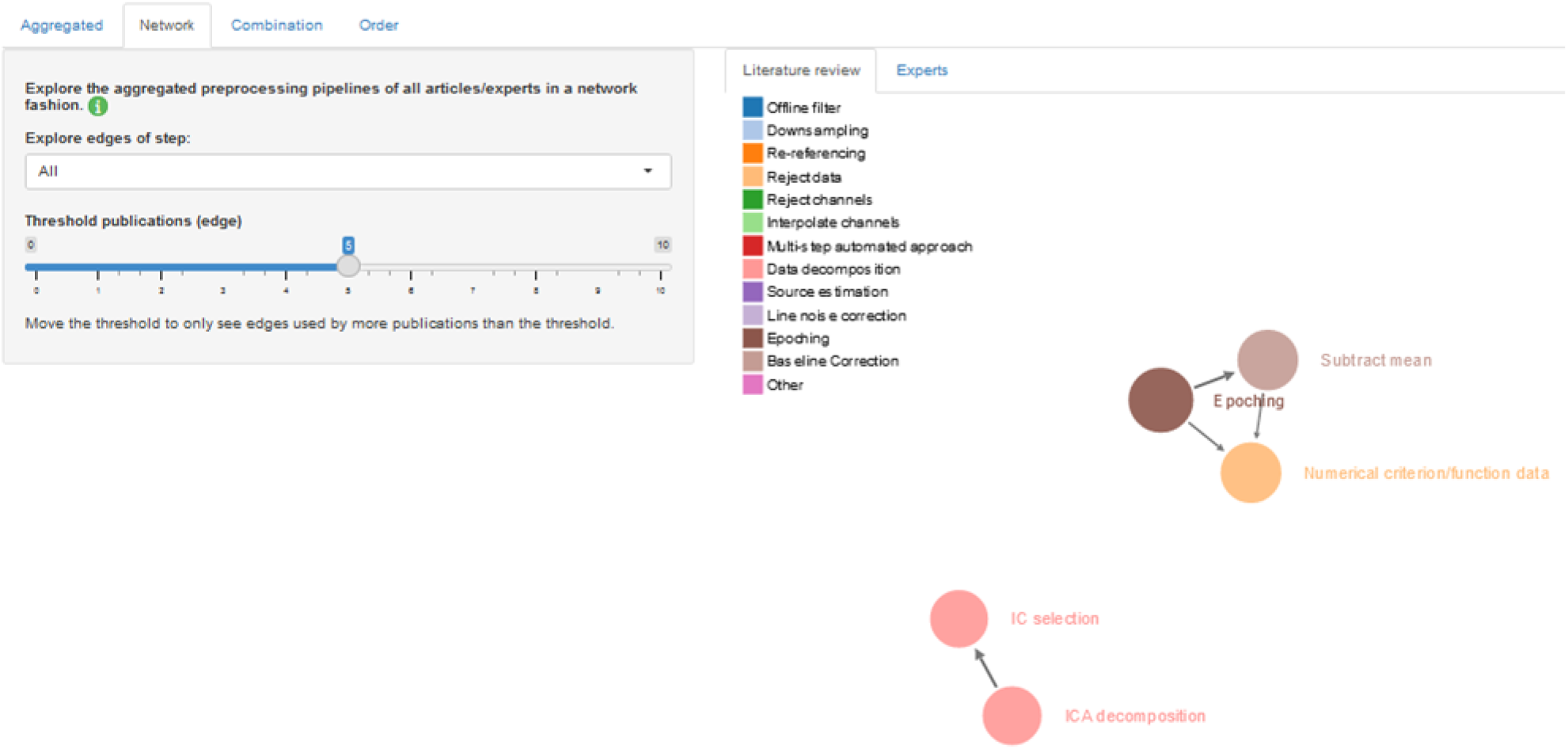
Edges of preprocessing steps as displayed in the Shiny app. Each node (circle) represents a preprocessing step. The color of a node indicates the processing group the step belongs to. Steps performed in succession are connected by arrows, these are edges. The wider the arrow, the higher the number of papers using this edge. The arrow points in the direction of the step that is performed afterward.

#### Which options of influential steps are common?

Prior studies have investigated the effect of several preprocessing choices on stationary EEG. Steps deemed influential are high-pass filtering (Clayson et al., 2021; Delorme, 2023; Šoškić et al., 2022), LPF (Clayson et al., 2021; Sadus et al., 2024; Schubert et al., 2023; Šoškić et al., 2022), re-referencing (Clayson et al., 2021; Delorme, 2023; Šoškić et al., 2022) and baseline correction (Clayson et al., 2021; Delorme, 2023; Šoškić et al., 2022)

HPF was performed by five studies with cut-off values ranging from 0.3 Hz to 1.5 Hz. LPF was used by 10 studies and the most used cut-off values were 40 Hz (*n* = 4) and 20 Hz (*n* = 3). Bandpass filtering (*n* = 17) was chosen more often than separate HPF and LPF. The lower cut-off frequencies ranged between 0.1 and 1 Hz and higher cut-offs between 30 to 100 Hz with the most popular passband being 1 to 30 Hz (*n* = 5). Most studies (n = 19) re-referenced the EEG data offline to a common average reference (CAR) (*n* = 12) or linked mastoids (*n* = 6) (see Figure *4*). A baseline correction was performed by 22 articles and a correction at -200 to 0 ms before stimulus onset was most common (*n* = 12).

**Figure 4.**
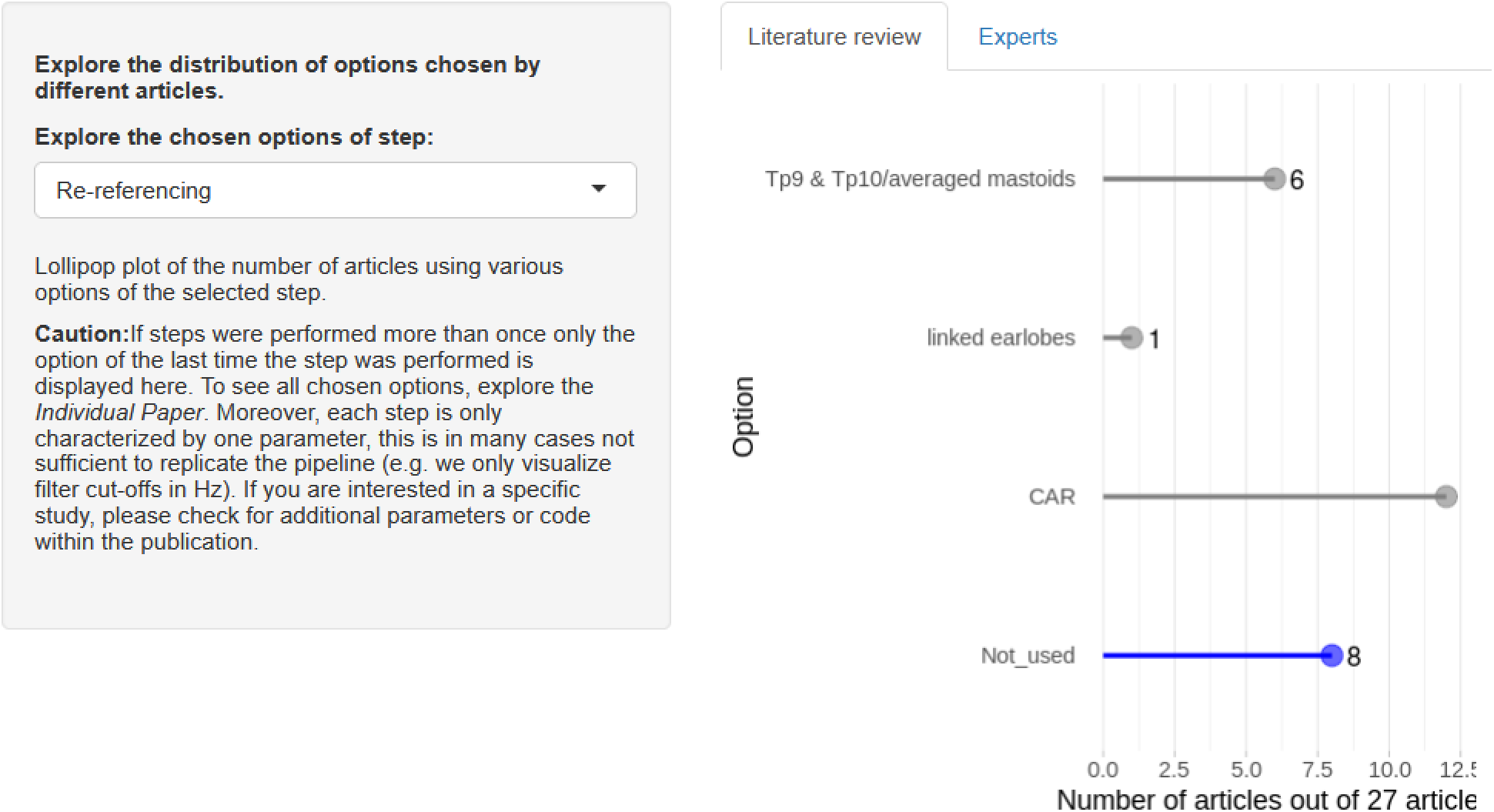
Options of the preprocessing step *Re-referencing* as displayed in the Shiny app. Number of articles using a specific option (grey) including common average re-referencing (CAR), the number of articles not using this step (blue), or are not reporting the chosen option (red, not displayed). Users can examine which articles chose which options within the Shiny app.

#### How could a preprocessing pipeline to investigate a P3 during walking look like?

Each article used a different preprocessing strategy to investigate a P3 during walking. We highlight an exemplary preprocessing from our lab, used in Scanlon et al. (2021b)(see Figure 5), to draw attention to principles used in many other pipelines. First, we recommend rejecting channels early, so they cannot affect artifact attenuation. Identifying channels with poor data quality is usually easier if it follows the removal of drifts and direct current offset with a high-pass filter. Secondly, ICA decomposition can help in attenuating stereotypical artifacts. It is important to note that interim preprocessing could be included to foster a good ICA decomposition. The resulting components can be applied to the original raw data, which can then be pre-processed appropriately for the feature of interest, such as the P3. ICA decomposition benefits from channel removal, a high-pass filter cut-off of 1 to 2 Hz (Debener et al., 2010; Klug & Gramann, 2020; Winkler et al., 2015), and the rejection of time points with abnormal values (Klug et al., 2024). One way of achieving this is generating 1s consecutive epochs and removing epochs with abnormal values based on joint probability or another numerical criterium. Various algorithms are available for ICA decomposition. Some may be more powerful than others (Delorme et al., 2012) and are commonly used in mobile EEG studies. ICs can be selected automatically to obtain replicable and scalable results. Yet reviewing automatic classification outcomes is strongly recommended since automatic procedures may not give valid results, especially when un-mixing results are poor. For instance, the neural and neck muscle activity of moving participants may not be separated well by ICA, making it difficult for automatic procedures to classify those components as representing brain or artifact signals. Thirdly, preprocessing should be appropriate for the feature of interest. Regarding filtering a P3 during motion, we had good experiences with a high-pass filter cut-off of 0.3 Hz, that removed some drifts but did not attenuate the P3 effect. In any case, the effects of temporal filtering on ERP morphologies must be very carefully evaluated (Widmann et al., 2015).

**Figure 5.**
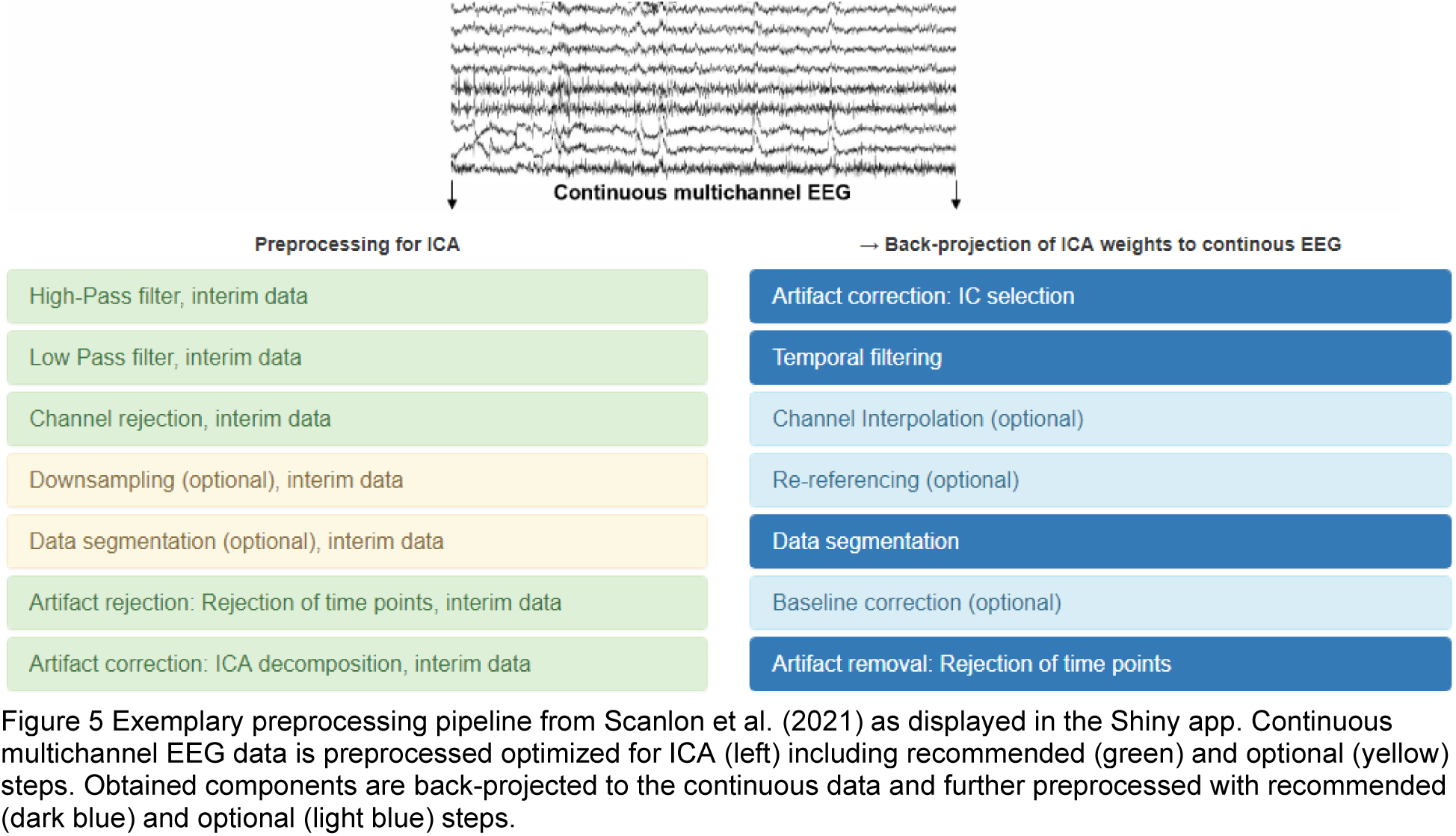
Exemplary preprocessing pipeline from Scanlon et al. (2021) as displayed in the Shiny app. Continuous multichannel EEG data is preprocessed optimized for ICA (left) including recommended (green) and optional (yellow) steps. Obtained components are back-projected to the continuous data and further preprocessed with recommended (dark blue) and optional (light blue) steps.

Note that this exemplary pipeline may serve as a starting point and does not necessarily represent the optimal preprocessing pipeline for a particular dataset. The pipeline is based on our experience with EEG signals captured during motion. Due to the influence of task, experimenter, subject, hardware, and contextual influences on EEG signal properties, other preprocessing pipelines may be adequate. In addition, this pipeline is no substitute for the empirical investigation of preprocessing choices on ERP results. For further reasoning on the order of steps, see the *exemplary pipeline* in the Shiny App.

#### Did anyone else perform the preprocessing pipeline I chose?

Users can construct their own preprocessing pipeline using the Shiny app by selecting the group and the step they want to perform. When clicking on a step, its definition is displayed, and – if available – a link to external resources for obtaining more information is provided. We also provide a primer on evaluating preprocessing pipelines (*info button > Download our primer*). Steps can be added with or without a specific option identified in the literature. Users can comment on new options, provide further information, such as version number or chosen parameter, or highlight if a step was performed on an interim dataset for ICA decomposition. Comments are not used when identifying similar pipelines. Users can download their preprocessing pipeline as a CSV file.

If one chooses a preprocessing pipeline that includes ICA decomposition, IC selection, epoching, and baseline correction (subtract mean) without choosing any specific option for these steps 15 studies are found that used the same steps. If the user searches for pipelines using these steps consecutively with the possibility of other steps in between, 10 studies are found. The exact same order of steps was used in three studies only (see Figure 6). Publications using the specified pipeline can be downloaded with the button on top of the table.

**Figure 6.**
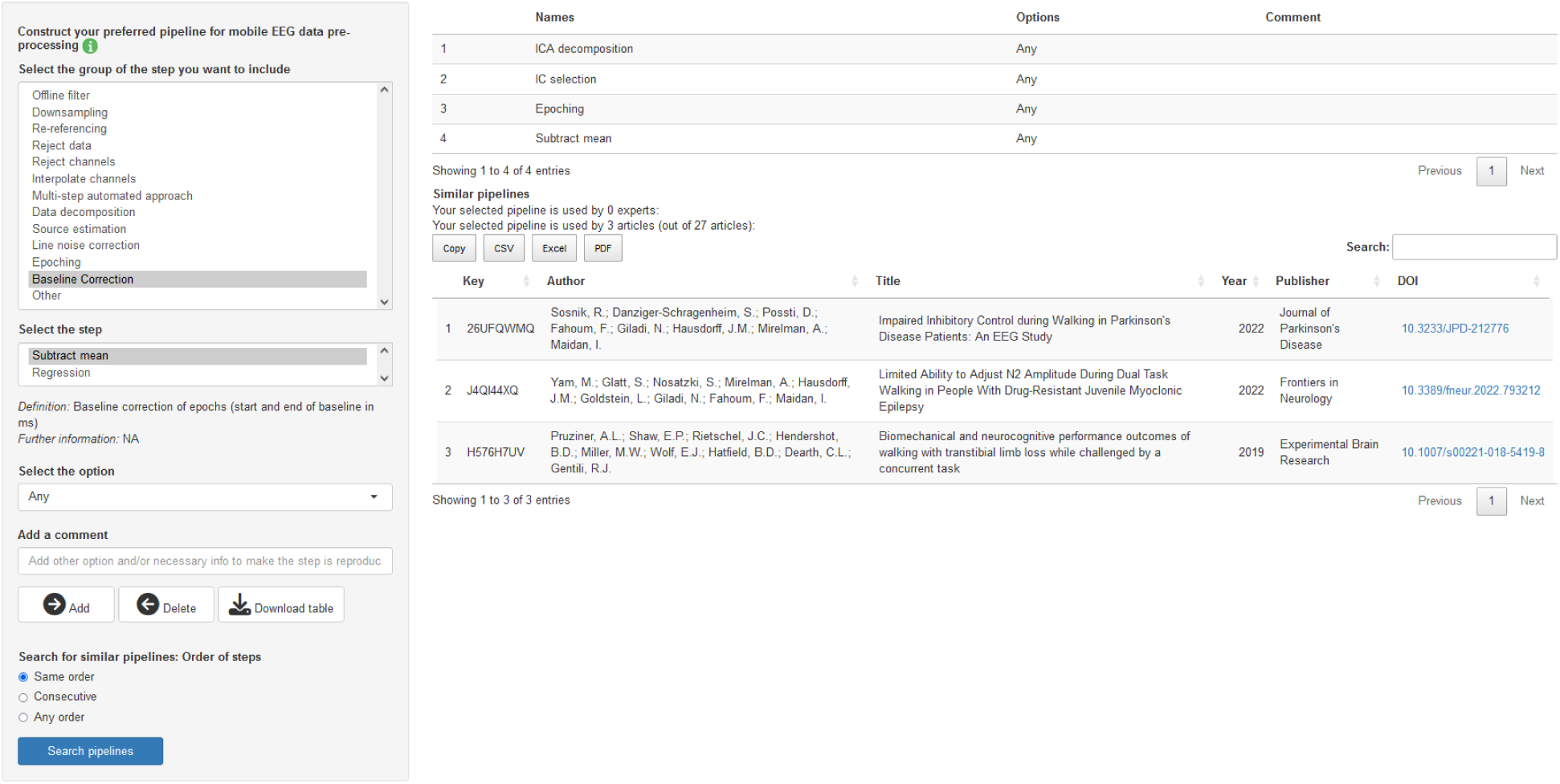
Constructed preprocessing pipeline as displayed in the shiny app. Steps and options can be selected and added. Using Search pipelines with the same steps and options can be identified.

## Discussion

We conducted a literature review of 27 studies on EEG preprocessing for P3 analyses in walking and stationary participants and could not identify a standard approach for analyzing mobile EEG. The number of employed preprocessing steps varied. Steps used by most authors include epoching, baseline correction, ICA, re-referencing, and bandpass filtering. Very few studies reported employing source estimation, line noise correction, or PCA-based methods. Previous reviews of mobile EEG point toward the potential pitfalls of consumer EEG systems which typically come with low channel counts (Gramann, 2024; Niso et al., 2022). Yet, most studies included in the present report used research-grade hardware with higher channel counts, which offers more flexibility with signal processing choices. Previous studies have examined the impact of some preprocessing choices on ERPs obtained from stationary, laboratory EEG recordings. In the following, we will compare the preprocessing choices we identified in the literature to recommendations from multiverse analyses of stationary ERPs. Delorme (2023) investigated the effect of single preprocessing choices to construct an optimized pipeline to detect experimental effects (i.e., the number of significant channels between conditions) in various datasets including an oddball P3. Clayson et al. (2021) examined preprocessing strategies aimed at enhancing data quality (i.e., between-trial standard deviations) and experimental effects (i.e., between-condition differences) the error positivity (Pe), and the error-related negativity (ERN). They provided an interactive visualization of the influence of preprocessing choices on the grand average ERP. Šoškić et al. (2022) explored the effect of preprocessing choices on the data quality (signal-to-noise ratio), effect size, and statistical power of a N400. For multiverse analyses concerning P3 latencies and their relation to cognitive abilities, refer to (Sadus et al., 2024; Schubert et al., 2023).

### Temporal filtering

In contrast to recommendations (Widmann et al., 2015), most studies used bandpass filtering instead of separating HPF and LPF with adjusted roll-off characteristics. Adequate HPF cut-off frequencies for ERP research have been debated and recent recommendations for P3 research range from 0.2 Hz (Zhang et al., 2024) to 0.5 Hz (Delorme, 2023) as ERP components can be altered by high HPF cut-offs (Duncan-Johnson & Donchin, 1979; Rousselet, 2012; Tanner et al., 2015). Some included studies surpassed these values. Six studies reported preprocessing an interim dataset, optimized for ICA decomposition. This may be due to optimized HPF settings for ICA. Especially for mobile studies, HPF cut-offs between 1 and 2 Hz are recommended to improve ICA decomposition (Debener et al., 2010; Klug & Gramann, 2020; Winkler et al., 2015). The detrimental effects of high filter cut-offs are usually circumvented by using higher HPF cot-offs before the ICA decomposition and back-projecting weights to a dataset with lower HPF filter cut-offs (Debener et al., 2010; Nenna et al., 2021; Protzak et al., 2021; Reiser et al., 2021; Scanlon et al., 2021a). In line with recent recommendations to use LPF cut-offs of 10 Hz or higher to investigate mean P3 amplitudes (Zhang et al., 2024), LPF cut-offs between 20 Hz and 40 Hz were most common in this review. LPF effects on the P3 were not investigated by Delorme (2023). LPFs with reasonable cut-offs of 30 Hz or higher did not affect mean amplitudes of a N400, while a low cut-off at 5.5 Hz smeared the peak in time (Šoškić et al., 2022). Comparing LPF cut-offs at 15, 20, and 30 Hz, Clayson et al. (2021) recommend 15 Hz LPF for ERN and 30 Hz LPF for Pe, reiterating that recommendations for one component may not hold for another – even within the same dataset. An interim dataset only used for ICA decomposition should be high-pass filtered with a cut-off between 1 to 2 Hz for better ICA decomposition. Filtering the data with separate high- and low-pass filters with cut-offs between 0.3 to 0.5 Hz and 30 to 40 Hz may work well for a P3 analysis, but we generally advise following well established ERP filter guidelines (Widmann et al., 2015).

### Re-referencing

In this review, most studies opted for offline re-referencing of EEG data, commonly employing average mastoids or CAR. Surprisingly, n No re-referencing strategy demonstrated an increase in experimental effect in a stationary oddball dataset; in fact, both average mastoids and CAR may even decrease an experimental effect (Delorme, 2023). Other investigations compared the impact of average mastoids versus CAR on various ERPs. For example, average mastoids were found to enhance the effect size and statistical power of the N400 compared to CAR (Šoškić et al., 2022). Conversely, Clayson et al. (2021) noted that CAR improved data quality, while average mastoids increased the experimental effect. Both linked mastoids and CAR are widely utilized re-referencing approaches, evident from their incorporation into several ERP multiverse analyses (Clayson et al., 2021; Delorme, 2023; Šoškić et al., 2022) and their utilization within mobile P3 literature. Unsurprisingly, all EEG reference and EEG re-reference challenges that are known for stationary EEG do also exist for mobile EEG and should not be ignored (Candia-Rivera et al., 2021; Delorme, 2023; Hagemann et al., 2001; Hu et al., 2019; Yao et al., 2019; Zheng et al., 2018).

### Baseline correction

Here, 22 out of 27 studies employed baseline correction, predominantly employing a pre-stimulus baseline ranging from -200 to 0 ms. Comparing pre-stimulus baselines of varying lengths, from 100 to 1000 ms to none in an oddball P3 paradigm, revealed that baseline correction did not augment experimental effects. Baseline durations of 500 ms or shorter were counterproductive for EEG data previously high-pass filtered at 0.5 Hz, although this observation may not extend to lower HPF settings (Delorme, 2023). In contrast, a separate investigation examining pre-stimulus baseline durations, from 100 to 350 ms, for the N400 component noted an increase in statistical power with longer baseline durations, provided that no confounding EEG activity occurred during the baseline period (Šoškić et al., 2022). In line with most reviewed studies, we typically use a baseline from -200 to 0 ms to obtain a stable estimate of the baseline activity.

While we could not identify a common strategy for P3 analysis, we describe common principles based on an exemplary preprocessing pipeline. These common principles include early rejection of channels and a separate interim dataset optimized for ICA decomposition with a high-pass filter of 1 to 2 Hz (Debener et al., 2010). Note that our experiences are no substitute for a proper empirical multiverse analysis that would be able to determine the impact of preprocessing choices on the statistical results.

Optimal preprocessing may also differ from study to study due to a variety of recording setups and employed tasks. Whether this should be reflected by tailored preprocessing, or we should strive for standardized preprocessing to investigate only the most robust effects is up to debate. In addition, optimal preprocessing may not only depend on the investigated ERP component (Clayson et al., 2021) but also the motion task as – for instance – slowly walking compared to running (Gwin et al., 2010; Nordin et al., 2019, 2020), playing table tennis (Studnicki et al., 2022) or even swimming (Klapprott & Debener, 2024) will give rise to different artifacts that artifact attenuation strategies may be sensitive and specific to, to varying extend (Jacobsen et al., 2022). In addition, mobile EEG, like stationary EEG, may be recorded with active pre-amplification at the electrode site or amplification only at the amplifier. In contrast to high-density stationary EEG, which is often used in research, mobile EEG systems vary considerably in the number of electrodes used, their spatial coverage, and the placement of the amplifier (Bateson et al., 2017; Gramann, 2024; Niso et al., 2022). These aspects of EEG acquisition may influence which preprocessing steps are most appropriate. For instance, Klug et al. (2024) observed that artifact rejection before ICA improved the decomposition -but to a much smaller extent than anticipated. Moreover, greater participant motion only reduced ICA decomposition quality within but not across different studies (Klug et al., 2024) highlighting the importance of assessing preprocessing impact with datasets similar to the data analyzed. Hence, adequate preprocessing of (mobile) EEG data may vary with the employed experimental task or EEG acquisition system. We propose evaluating the impact of a limited number of preprocessing decisions on a dataset with similar EEG data characteristics, such as a pilot dataset or prior work within the lab. One can include common preprocessing choices in P3 analysis which can be identified with the shiny app. Moreover, recommendations can help identify suitable and preprocessing approaches, as well as set-up procedures and EEG systems (Gorjan et al., 2022; Gramann, 2024; Reis et al., 2014; Song & Nordin, 2021; Wascher et al., 2021). If a scaffolding of steps exists, employing a multiverse analysis using data that will not undergo statistical evaluation can determine appropriate options for specific steps.

Given that no standards for EEG P3 preprocessing exist and that limited resources may prevent researchers from conducting extensive multiverse analysis, we should ensure that preprocessing is at least reproducible. Unfortunately, EEG analysis details are not always reported in sufficient detail (Gramann, 2024; Šoškić et al., 2021). Two possible approaches to tackle this problem may be considered. First, authors should be encouraged to adhere to existing reporting standards. Second, authors should be encouraged to embrace open scholarship and share the code for data processing and the processed data. Among the existing reporting standards for EEG, we wish to highlight two. First, the *Best Practices in Data Analysis & Sharing in Neuroimaging using MEEG* (COBIDAS-MEEG) (Pernet et al., 2020) provide good practices for magnetoencephalography and EEG (MEEG) data acquisition, analysis, and reporting. A living white paper with the guidelines can be found at https://cobidasmeeg.wordpress.com/. Second, the ARTEM-IS (Šoškić et al., 2023; Styles et al., 2021) provides reporting standards for ERP research. A web app (available at https://artemis.incf.org/) allows authors to create a template to report all methodology details of a study. This report can be shared and/or downloaded as PDF or JSON file. Both guidelines have a scope beyond EEG preprocessing encompassing EEG acquisition, parametrization, and statistical evaluation. They may, however, require adaptations to address challenges in mobile EEG research. It may be argued that methodological descriptions are too excessive already. This problem could be easily addressed by including extended methods sections in supplemental materials or sharing them in a respective repository. An increasing number of mobile EEG studies embrace the potential of recording brain activity in ecological settings. While this may enhance ecological validity (depending on the context and used task), it compromises reproducibility due to uncontrollable environmental variations among participants (Holleman et al., 2020). Therefore, meticulous documentation of recording contexts is imperative.

Apart from using existing standards, code should be shared since documentation and implementation may deviate. Moreover, shared code is crucial if manual steps (e.g., visual inspection to identify noisy data segments or artifactual ICs) are employed to achieve a reproducible workflow. Open scholarship practices were not as widely employed as we hoped although 81% of the studies were published following the replication crisis (Open Science Collaboration, 2015) which fostered their use. Interestingly, open data was the most used practice, although the sharing of human neuroimaging data is (in contrast to the sharing of code and material) subject to privacy concerns (Gaspar et al., 2011; Kong et al., 2019; Lai et al., 2022; Ruiz-Blondet et al., 2017). The two approaches may be best combined.

## Outlook

This literature review and the accompanying Shiny app are a step toward the identification of the mobile EEG multiverse. To not obtain biased results or overlook impactful choices, only defensible decisions should be included in a multiverse analysis (Del Giudice & Gangestad, 2021). Here, only preprocessing strategies reported in peer-reviewed publications were examined. To consolidate that these are defensible and to extend to novel preprocessing strategies, mobile EEG experts could be surveyed and/or preprocessing pipelines may be discussed in focus groups to see which decisions are justified among a larger group of experienced users. After defensible forking paths are established and evaluation metrics are defined, multiverse analysis may be used to identify the impact of different preprocessing strategies on EEG data. Likely, which preprocessing strategy is well suited varies across datasets and may depend on parameters of the EEG acquisition system (e.g., amplifier weight and placement). This could be evaluated with simulated data, for which the signal of interest is known (Richer et al., 2019, 2020).

While we did not expect to identify a standard for P3 preprocessing in mobile EEG studies during walking, certain processing steps were widely employed and may help to foster consensus. To advance the field, quality metrics may be used to assess EEG data quality and to identify optimal preprocessing pipelines within the multiverse. Still, expert assessments are crucial to ensure that only defensible forking paths are evaluated. In the meantime, authors should take great care to provide detailed metadata on EEG acquisition, preprocessing, and parametrization, so that results may be replicated using existing reporting standards. Furthermore, we encourage authors to share documented and structured code openly. This not only facilitates replication but also serves as an additional layer of documentation, contributing to the overall transparency and integrity of the research process.

This review provides a snapshot of preprocessing choices of one ERP component, the P3, during walking that were published at the time of writing. More studies investigating this component will be conducted in the years to come. To keep the app up to date a semi-automatic extraction framework, fine-tuned with steps and options identified in the present review, could identify suitable articles and extract information automatically akin to the automatic extraction of functional magnetic resonance imaging coordinates in NeuroSynth (RRID:SCR_006798).

## Limitations

This review only investigated the preprocessing choices of 27 studies. Our ability to identify commonalities between studies may have increased with a larger sample size which may be available in the future. Moreover, we only focused on one ERP component, namely the P3. The use of automated tools for information extraction (see Outlook) could have allowed us to investigate more studies, also including other EEG measures. In addition, we only coded one option for each step. For instance, we only extracted the filter cut-off value in Hz for LPFs. Yet, filters are characterized by various other features, such as impulse response time, windowing function, filter order, causality, and phase shift which should be reported as well (Pernet et al., 2020). Hence, our finding of all studies reporting their filter cut-offs in Hz does not indicate that these filters are reported in sufficient detail to be reproduced. Furthermore, the Shiny app does not allow to discriminate if preprocessing steps were only conducted on an interim dataset. For example, 16 studies reported using a high-pass or bandpass filter with a frequency cut-off of 1 Hz or higher which may influence the resulting ERP (Tanner et al., 2015) but may have been performed on interim datasets for optimized ICA performance and weights could have been back-projected to a dataset with a much lower HPF suited for ERP analyses. This information is not well represented here but can be looked up in the original publications. Finally, our exemplary use case focused on finding out which practices are common in the examined part of the literature, but we did not assess the impact of these preprocessing steps. Nonetheless, common decisions in peer-reviewed publications should reflect an acceptance by the community.

## Conclusion

In summary, our exploration of EEG preprocessing for mobile P3 analyses highlights the need for standardized practices, thorough reporting, and a commitment to open science. The proposed app allows interactive exploration of the database. By addressing these considerations, the scientific community can advance the reliability and reproducibility of mobile P3 studies, fostering a more robust foundation for future research in this domain.

## Data and Code Availability

The data and code supporting the findings of this study, as well as the source code of the shiny app, are available at https://github.com/metascience-uol/mobileEEG_multiverse. The repository also provides a link to the app, and a Docker image that can be accessed at (docker pull nadinejacobsen/meteor:eeg_meteor). The image then can be run locally using the command docker run –p 3838:3838 nadinejacobsen/meteor:eeg_meteor. The app will then be accessible at http://localhost:3838.

## Author Contributions

**NJ**: Conceptualization, Methodology, Software, Data Curation, Writing - Original Draft. **DK**: Software, Writing - Review & Editing. **SW:** Investigation, Data Curation, Writing - Review & Editing. **CI**: Software, Writing - Review & Editing. **SD:** Conceptualization, Writing - Review & Editing, Supervision

## Footnotes

1 Since some studies compared several systems, we report results from 32 systems used in 27 studies.

